# Analysis of differentially methylated regions in primates and non-primates provides support for the evolutionary hypothesis of schizophrenia

**DOI:** 10.1101/322693

**Authors:** Niladri Banerjee, Tatiana Polushina, Francesco Bettella, Vidar M. Steen, Ole A. Andreassen, Stephanie Le Hellard

## Abstract

**Introduction:** The persistence of schizophrenia in human populations separated by geography and time led to the evolutionary hypothesis that proposes schizophrenia as a by-product of the higher cognitive abilities of modern humans. To explore this hypothesis, we used here an evolutionary epigenetics approach building on differentially methylated regions (DMRs) of the genome.

**Methods:** We implemented a polygenic enrichment testing pipeline using the summary statistics of genome-wide association studies (GWAS) of schizophrenia and 12 other phenotypes. We investigated the enrichment of association of these traits across genomic regions with variable methylation between modern humans and great apes (orangutans, chimpanzees and gorillas; primate DMRs) and between modern humans and recently extinct hominids (Neanderthals and Denisovans; non-primate DMRs).

**Results:** Regions that are hypo-methylated in humans compared to great apes show enrichment of association with schizophrenia only if the major histocompatibility complex (MHC) region is included. With the MHC region removed from the analysis, only a modest enrichment for SNPs of low effect persists. The INRICH pipeline confirms this finding after rigorous permutation and bootstrapping procedures.

**Conclusion:** The analyses of regions with differential methylation changes in humans and great apes do not provide compelling evidence of enrichment of association with schizophrenia, in contrast to our previous findings on more recent methylation differences between modern humans, Neanderthals and Denisovans. Our results further support the evolutionary hypothesis of schizophrenia and indicate that the origin of some of the genetic susceptibility factors of schizophrenia may lie in recent human evolution.

## 1. Introduction

Schizophrenia is a psychiatric disorder with a prevalence rate of 2.7-8.3/1,000 persons (Messias et al., 2007) and heritability estimated between 60-90% (Cardno et al., 1999; Lichtenstein et al., 2009; Skre et al., 1993; Sullivan et al., 2003). It occurs at quite similar rates across populations worldwide (Ayuso-Mateos, 2002; Brüne, 2004; WHO, 1973) and written records describing its symptoms exist dating back 5,000 years (Jeste et al., 1985). This consistent persistence of the disease despite reduced fecundity (Brüne, 2004; Nichols, 2009) and increased mortality is a paradox (Bassett et al., 1996; Brown, 1997; Larson and Nyman, 1973), since the reduced fecundity of patients afflicted with schizophrenia does not appear to eliminate the disease from the population (Power et al., 2013) Part of the reason may be due to afflicted individuals reproducing prior to the onset of the disease (Markow, 2012). Another contributing factor could be that schizophrenia risk variants may have provided an advantage to the kin of the affected by conferring superior creative and intellectual abilities upon them (Kyaga et al., 2011; Nichols, 2009). To explain the constant occurrence of the disease, TJ Crow (Crow, 1997, 1995) proposed the so-called evolutionary hypothesis of schizophrenia, which suggests that the disease is a consequence of human evolution: the higher cognitive abilities of modern-day humans, including language, may predispose to psychiatric illnesses such as schizophrenia (Crow, 2008, 2000, 1997).

In the post-genomic era (Lander et al., 2001; Venter et al., 2001), emerging lines of evidence are lending support to this hypothesis. Crespi et al. (Crespi et al., 2007) were amongst the first to show that genes with evidence of recent positive selection in humans are also implicated more frequently in schizophrenia. More evidence has been provided by studies based on comparative genomics (Pollard et al., 2006; Srinivasan et al., 2015; Xu et al., 2015), a field in which genomes of progressively older species are compared to identify substitutions and mutations that help estimate divergence between the species. For instance, a group of regions defined by negative Neanderthal selective sweep (NSS) scores describe the selective evolution of genomic regions in modern-day humans over Neanderthals (Burbano et al., 2010; Green et al., 2010). These regions were shown by Srinivasan et al. (2015) to be enriched for schizophrenia risk markers, in line with the evolutionary hypothesis of schizophrenia. Other regions known as human accelerated regions (HARs) (Gittelman et al., 2015; Pollard et al., 2006; Xu et al., 2015), first described by Pollard et al. (2006), show accelerated evolution in humans compared to primates or mammals. HARs have also provided some evidence of enrichment of association with schizophrenia (Xu et al., 2015), but these findings may have been driven by a few genes since they were not replicated using a polygenic approach (Srinivasan et al., 2017, 2015).

While several studies have looked at the evolution of the genome (Bird et al., 2007; Bush and Lahn, 2008; Gittelman et al., 2015; Paaby and Rockman, 2014; Pollard et al., 2006), there are reports that the epigenome is evolving as well (Gokhman et al., 2014; Hernando-Herraez et al., 2015, 2013; Mendizabal et al., 2014; Molaro et al., 2011). This provides new insights into events leading to the speciation and divergence of modern humans. The epigenome refers to the layer of chemical modifications, such as methylation and histone modifications, to the genome that regulate gene expression (Bernstein et al., 2007; Kundaje et al., 2015; Rivera and Ren, 2013). For instance, Gokhman et al. (2014) compared the methylomes of humans with Neanderthals and Denisovans. They reported that while 97% of the methylome was comparable between humans, Neanderthals and Denisovans, some regions showed differential methylation between the three hominids. Previously (Banerjee et al., 2017), we analysed the differentially methylated regions (DMRs) identified for Neanderthals, Denisovans and modern humans by Gokhman et al. (2014), and found evidence that the regions of the genome with human-specific DMRs harbour relatively more genetic variants associated with schizophrenia than the rest of the genome, i.e. the DMRs were enriched for SCZ markers both at the single-nucleotide polymorphism (SNP) level and at the gene level. These human-specific DMRs thus provide evidence of enrichment of methylation changes in regions harbouring genetic variants associated with schizophrenia, at least since the divergence from Neanderthals and Denisovans (Banerjee et al., 2017).

Here, we sought to determine if evolutionarily older methylation differences can provide a further timeframe for the origin of schizophrenia risk markers in the human lineage. We asked whether we can find epigenetic evidence that the origin of schizophrenia risk markers predates the origins of the Homo genus, i.e. before the divergence of chimpanzees and humans around 6-8 million years ago (MYA) (Glazko and Nei, 2003; Langergraber et al., 2012). We tested this hypothesis by analysing primate DMRs that trace an evolutionary history of at least 13 million years (Glazko and Nei, 2003; Hasegawa et al., 1985; Rannala and Yang, 2003). We used the same statistical analyses as described by Lee et al. (2012), Schork et al. (2013), and Srinivasan et al. (2015) to test for polygenic enrichment of a set of markers from genome-wide association studies (GWAS). We interrogated regions of the human genome which are hypo- or hyper-methylated in comparison to the corresponding ones in chimpanzees, gorillas and orangutans for enrichment of genetic variants associated with schizophrenia or other human traits.

## 2. Materials and methods

### 2.1. GWAS data

Summary statistics for thirteen different phenotypes were obtained from their respective published GWAS studies: schizophrenia (SCZ) (Ripke et al., 2014), bipolar disorder (BPD) (Sklar et al., 2011), attention deficit hyperactivity disorder (ADHD) (Demontis et al., 2017), rheumatoid arthritis (RA) (Stahl et al., 2010), blood lipid markers (high density lipoprotein (HDL), low density lipoprotein (LDL), triglycerides (TG), total cholesterol (TC)) (Teslovich et al., 2010), blood pressure (systolic blood pressure (SBP), diastolic blood pressure (DBP)) (Ehret et al., 2011), body mass index (BMI) (Locke et al., 2015), height (Wood et al., 2014) and intelligence (Sniekers et al., 2017). For studies published with hg18 coordinates (BPD, SBP, DBP, HDL, LDL, TG, TC, RA), conversion to hg19 was performed using the command line version of the liftOver tool from the UCSC Genome Browser (Karolchik et al., 2014) (http://hgdownload.cse.ucsc.edu/downloads.html#utilities_downloads). For BMI and height SNPs, the genomic coordinates were obtained by mapping them to the assembly of 1,000 Genomes Project (1KGP) Phase 1 reference panel SNPs (Durbin et al., 2012).

### 2.2. Human hypo- and hyper-methylated regions from primate DMRs

These methylated regions were retrieved from the study by Hernando-Herraez et al. (2015), who identified them by comparing the methylation profile of DNA from peripheral blood samples of orangutans, chimpanzees and gorillas to that of humans. Since the DMRs are determined by comparing humans with other primates, we refer to this set collectively as primate DMRs. Both hypo- and hyper-methylated DMRs from humans were analysed. As these DMRs are identified in the 175 same tissues in all samples, they are considered to represent species-specific methylation differences, not tissue-specific methylation differences (Gokhman et al., 2014). Altogether, the human hypo- and hyper-methylated DMRs can be used to represent an evolutionary course of history spanning from at least 13 MYA (Glazko and Nei, 2003; Langergraber et al., 2012), when orangutans diverged from the common ancestors, to 6 MYA, when the chimpanzees and humans diverged from each other (Glazko and Nei, 2003; Langergraber et al., 2012). Since our interest was in human-specific enrichment, we focused the analyses on human hypo- and hyper-methylated DMRs.

### 2.3. Differentially methylated regions (DMRs) from Neanderthals, Denisovans and modern humans

As previously described (Gokhman et al., 2014), these methylated regions have been identified by comparing the methylomes of osteoblasts from modern-day humans with those from Neanderthals and Denisovans. We refer to them in this paper as non-primate DMRs. Gokhman et al. (2014) devised a strategy utilizing information in the form of cytosine (C) to thymine (T) ratios to decipher the ancient methylomes of Neanderthals and Denisovans. Subsequently, they compared the methylomes of Neanderthals, Denisovans and modern humans and inferred the species in which the methylation variation likely took place; this information was used to classify the DMRs as Neanderthal-specific, Denisovan-specific and human-specific. These DMRs represent species-specific methylation (Gokhman et al., 2014).

### 2.4. Neanderthal selective sweep (NSS) data

We obtained NSS marker data from Srinivasan et al. (2015). Negative scores for NSS markers indicate positive selection in humans. Markers with such scores were used in the downstream analyses.

### 2.5. SNP assignment with LDsnpR

The previously published R-based software package LDsnpR (Christoforou et al., 2012) was utilized for assigning SNPs to the respective DMRs using LD (linkage disequilibrium)-based binning at r^2^ ≥0.8 in R (R Core Team, 2017). LD-based binning makes it possible to determine whether SNPs from a specific GWAS are in LD with the DMR of interest. Using LD allows the capture of a greater number of relevant SNPs in comparison to an approach where only physically overlapping SNPs are considered. The LD file utilized was in HDF5 format and was constructed from the European reference population of 1KGP and can be publicly downloaded at: http://services.cbu.uib.no/software/ldsnpr/Download.

### 2.6. Enrichment analyses based on stratified quantile-quantile (QQ) plots

QQ plots are an essential method used in GWASs to depict the presence of true signals. They help to visually observe the spread of data and deviations from the null distribution. Under the null hypothesis, no difference is expected between the observed and expected distributions of data. As such, a line of no difference or null line is obtained that is equidistant from both X and Y axes. However, if the null hypothesis were to be false, there would be a deviation of the observed data distribution from the expected data distribution. As described in depth by Schork et al. (2013), a leftward deflection of the observed distribution from the null line represents enrichment – the greater the leftward deflection, the stronger the enrichment of true signals. This method has been used recently not only to show how specific genomic annotation affects the distribution of disease SNPs with true signals (Schork et al., 2013), but also to demonstrate that regions of recent evolution are enriched for schizophrenia markers (Banerjee et al., 2017; Srinivasan et al., 2015). We took the SNPs that are in LD with the DMR regions and plotted their *p*-value distributions from various GWASs. The observed *p*-value distributions were then determined to be enriched or not using conditional Q-Q plots as described by Schork et al. (2013). Genomic inflation was corrected by *λ*_GC_.

### 2.7. INRICH-based enrichment analysis

The stratified QQ plots provide a visual depiction of data distributions and enrichment of true signals within a stratum of data, but they do not quantify this enrichment. Therefore, we used the INterval EnRICHment (INRICH) analysis tool to statistically quantify the enrichment observed. This pipeline performs permutation and bootstrapping procedures to determine with statistical confidence whether LD-implicated genomic intervals are enriched in specific gene sets (Lee et al., 2012). The INRICH analysis takes into account several potential biases that can otherwise lead to false positives, such as variable gene size, SNP density within genes, LD between and within genes, and overlapping genes in the gene sets. We used the same procedure reported previously (Banerjee et al., 2017; Xu et al., 2015) with SNPs in the extended MHC region and SNPs with MAF <0.05 excluded from the analysis. Additional details can be found in the Supplementary Information.

## 3. Results

### 3.1. Co-localisation of human hypo-methylated regions and genetic variants associated with schizophrenia in the MHC

We ascertained whether there is any enrichment of human hypo- and hyper-methylated regions in schizophrenia-associated SNPs. Using previously published methodology (Christoforou et al., 2012), we mapped schizophrenia markers to human hypo-methylated regions (hypo-DMRs) and hyper-methylated regions (hyper-DMRs). Out of a total of ~9.4 million SCZ markers obtained from the GWAS, 10,165 markers tagged hypo-DMRs and 4,503 tagged hyper-DMRs.

Figure 1A shows the conditional QQ plots for schizophrenia markers (all markers, the hypo-DMR set and the hyper-DMR set) including those in the MHC region. For hypo-DMR markers (Supplementary Dataset 1), we observed a significant enrichment as depicted by the leftward deviation. No enrichment was observed for hyper-DMR markers. Since the MHC region is a region of extended linkage disequilibrium, which can bias the enrichment estimates, and since it is the main region of association with schizophrenia, we also tested the enrichment with the MHC region removed (Figure 1B). Under these conditions there is a trend for enrichment of hypo-DMR markers at higher *p*-value thresholds, but this enrichment is substantially less than when the MHC is included (Figure 1A).

**Fig. 1:**
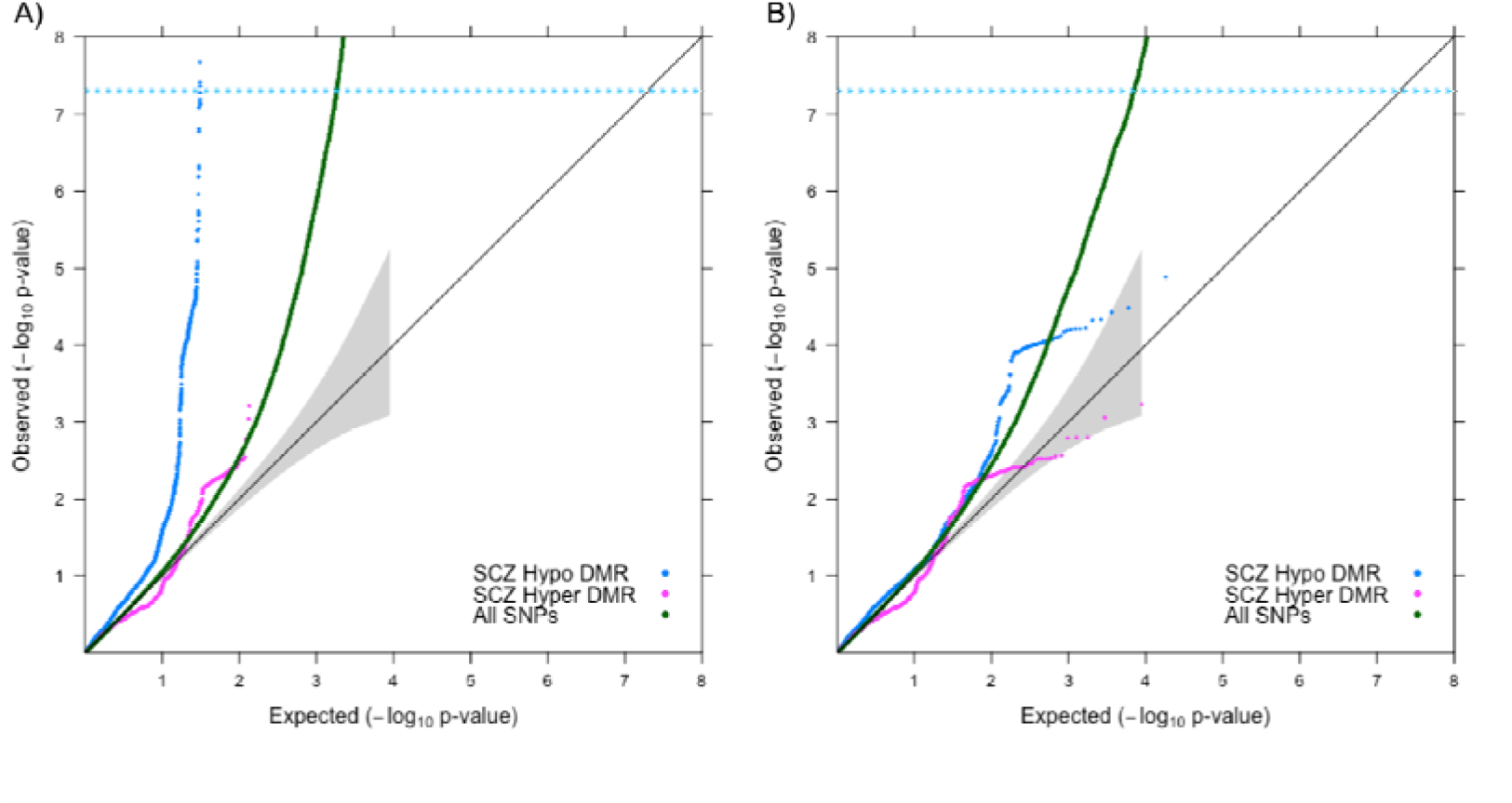
Enrichment plots of hypo-DMR and hyper-DMR SNPs in schizophrenia. Quantile-quantile (QQ) plots of GWAS SNPs for Schizophrenia (SCZ) with the extended MHC region (chr6: 25-35Mb) unmasked (A) and masked (B). Expected - log_10_ *p*-values under the null hypothesis are shown on the X-axis. Observed -log_10_ *p*-values are on the Y-axis. The values for all GWAS SNPs are plotted in dark green while the values for SNPs in linkage disequilibrium (LD) with hypo-methylated DMRs are plotted in blue and SNPs in LD with hyper-methylated DMRs are plotted in pink. A leftward deflection of the plotted *p*-values from the line for all GWAS SNPs indicates enrichment of true signals – the greater the leftward deflection, the stronger the enrichment. Genomic correction was performed on all SNPs with global lambda.

### 3.2. Enrichment of markers is not seen for other human traits

Next, we tested if the human hypo- and hyper-methylated regions are enriched for other human traits and phenotypes. We tested a total of thirteen different phenotypes, full details of which can be found in section 2.1. Each GWAS had been performed with a different number of genotyped SNPs, and this difference could potentially bias our results. To circumvent this, we created a list of ~2.4 million common SNPs that were genotyped across all the phenotypes investigated in the present study. Only SNPs on this list were used for enrichment analysis.

As can be seen in Fig. 2, no enrichment was observed in any of the traits, with the possible exception of height at higher *p*-value threshold markers. The common list of markers did not contain the MHC region and as such no enrichment is observed for schizophrenia either.

**Fig. 2:**
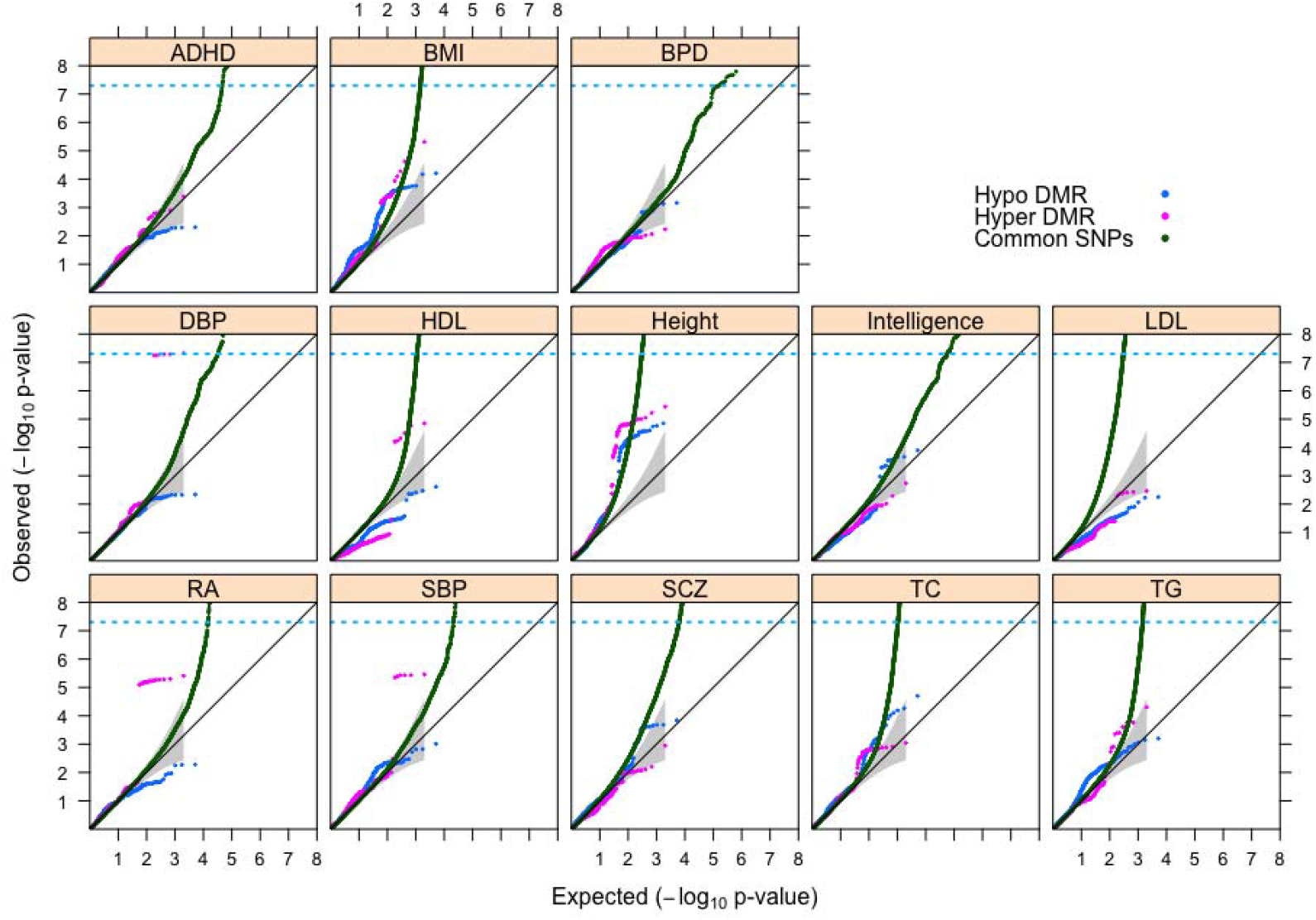
Enrichment plots of hypo-DMR and hyper-DMR SNPs across multiple traits. Thirteen different GWASs were analysed using a common set of ~2.4 million SNPs. The *p*-values for the common set of GWAS SNPs are plotted in dark green; *p*-values for SNPs that tag hypo-methylated DMRs are plotted in blue; and *p*-values for SNPs that tag hyper-methylated DMRs are plotted in pink. ADHD, attention deficit hyperactivity disorder; BMI, body mass index; BPD, bipolar disorder; DBP, diastolic blood pressure; HDL, high density lipoprotein; LDL, low density lipoprotein; RA, rheumatoid arthritis; SBP, systolic blood pressure; SCZ, schizophrenia; TC, total cholesterol; TG, triglycerides. The MHC region was absent from the common set of SNPs.

### 3.3. Evidence of enrichment for hypo-methylated regions with SNPs at high p-values

The enrichment plots allowed us to visually ascertain enrichment in the datasets. However, they did not give any indication of the statistical robustness of the enrichment. To ascertain if the human hypo- and hyper-methylated regions are statistically enriched for schizophrenia and height markers, we implemented the INRICH pipeline, which performs 10,000 permutations and 5,000 bootstrapping calculations, to determine with statistical confidence the enrichment observed (Lee et al., 2012).

The INRICH analysis confirmed a significant (*p<*0.05) enrichment of association for human hypo-DMRs, but not hyper-DMRs, with schizophrenia at SNPs of higher *p*-value thresholds *(p*<10e-3 to *p*<10e-4) (Fig. 3). This enrichment was at the gene level, and complemented the enrichment observed at the SNP level for higher *p*-value thresholds (Fig. 1B). Importantly, this enrichment persisted upon testing a pruned schizophrenia dataset (Supplementary Fig. 1). The enrichment was however not significant at the genome-wide threshold (*p<*5×10e-8) and was much weaker than that observed for non-primate DMRs (Fig. 3). We also observed a similar trend for height where there was enrichment at SNPs of higher but not lower *p*-value thresholds. This enrichment was similarly less pronounced than for non-primate DMRs (Supplementary Fig. 2).

**Fig. 3:**
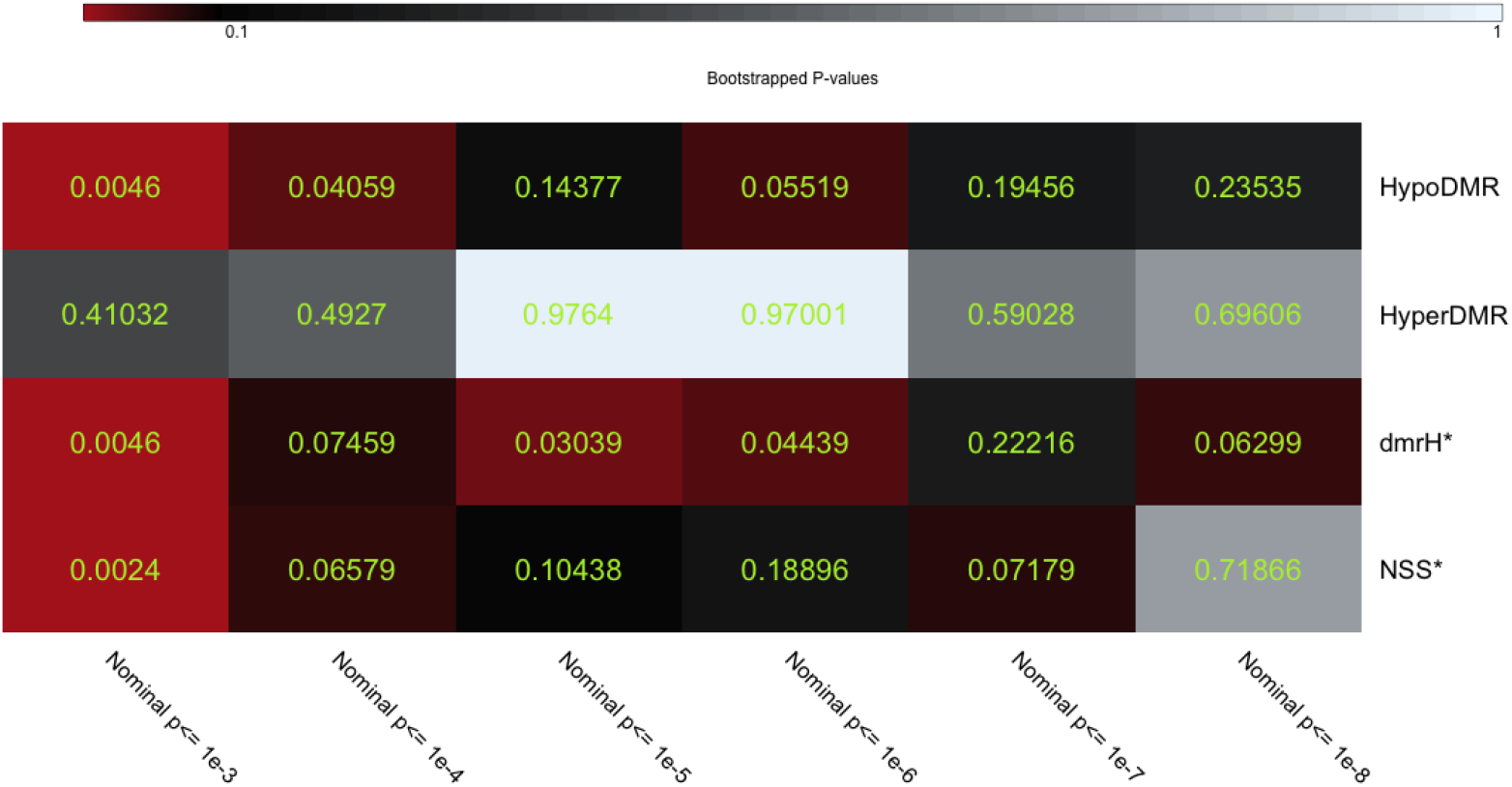
INRICH test for enrichment of association of DMR gene sets and NSS genes with SCZ, MHC masked. A visual heatmap depicting *p*-values from bootstrapping with 5,000 iterations. The various evolutionary annotations compared are as follows. HypoDMR – human hypo-methylated DMRs; HyperDMR – human hyper-methylated DMRs. HypoDMR and HyperDMR were taken from the study by Hernando-Herraez et al. (2013). dmrH – human-specific DMRs (Gokhman et al, 2014), which are referred to as non-primate DMRs in this manuscript. NSS - Neanderthal selective sweep. Datasets marked with * have been previously reported by Banerjee et al. (2017) and are presented here for comparison only.

## 4. Discussion

In our study, we investigated if regions of the human genome whose methylation has evolved since the divergence of modern humans from great apes are enriched for markers of schizophrenia. We found evidence that there is enrichment for hypo-methylated DMRs driven by the MHC locus, a known risk region that harbours the most significant schizophrenia GWAS markers (Ripke et al., 2014). When the MHC region was excluded from the analysis, there remained a trend towards enrichment of hypo-DMRs driven by SNPs of higher *p*-value thresholds. This finding was complemented by the INRICH analyses that indicated significant enrichment among SNPs of higher *p*-value thresholds. When analysing a global SNP list common to GWAS of several traits, we failed to find evidence of enrichment of any trait with the possible exception of height at higher SNP *p*-value thresholds. We tested this further with the INRICH pipeline, which revealed gene-level enrichment of LD intervals for height markers below the genome-wide threshold (*p*<5×10e-8). Compared to our previous study, in which we demonstrated enrichment of association with schizophrenia for non-primate DMRs that were derived by comparing human, Neanderthal and Denisovan methylomes (Banerjee et al., 2017), the primate DMRs tested here show far less enrichment. The primate and non-primate DMRs have very little overlap, which suggests that the methylation changes that took place since the divergence of modern humans from Neanderthals and Denisovans occurred in different regions of the genome compared to those that took place since divergence from great apes.

The central role of the MHC region in the enrichment of human hypo-methylated regions poses interesting questions. The MHC region is known for its complex LD architecture, which renders the interpretation of genetic signals very challenging. Other groups have previously reported that the MHC region is one of the fastest evolving regions of the human genome (Meyer et al., 2017) and have implicated it in mate preference (Bernatchez and Landry, 2003; Kromer et al., 2016; Potts and Wakeland, 1990; Roberts et al., 2008; Winternitz et al., 2017), odour perception (Roberts et al., 2008; Santos et al., 2005) and immune response (Benacerraf, 1981; Horton et al., 2004). Recently it was shown that a large proportion of the association of the region with schizophrenia can be explained by complement C4 haplotypes that include C4 copy number variation (Sekar et al., 2016). Nevertheless, there remains a part of the association in this region that is unexplained (Gejman et al., 2011) and will need further investigation. It is interesting to consider the possibility that the MHC region and the immune system in general play a central role in evolution at the epigenomic as well as at the genomic level (Meyer et al., 2017; Potts and Wakeland, 1990; Sommer, 2005; Traherne, 2008). The mechanisms by which hypo-methylation could influence the aforementioned processes are open to speculation since the MHC region has more than 200 genes in close physical proximity and LD with one another (Beck et al., 1999). This makes it hard to interpret the exact biological consequences of our findings.

Interestingly, the gene-level analysis via INRICH seems to suggest enrichment of SNPs of higher *p*-value thresholds in primate DMRs for both schizophrenia and height. This enrichment is far lower than what we found for non-primate DMRs for both schizophrenia and height (Banerjee et al., 2017) and which persisted for schizophrenia even with pruned datasets.

The very small overlap between primate and non-primate DMRs might suggest that the divergence from Neanderthals and Denisovans brought about more significant methylation changes in regions implicated in the aetiology of schizophrenia and height than the divergence from great apes. In other words, our results might suggest that the evolutionary factors that regulate methylation variation acted on different segments of the genome at different time points. So while the methylation variation since the divergence from Neanderthals and Denisovans may mark a genome-wide increase of schizophrenia susceptibility (Banerjee et al., 2017), the methylation variation from the time period between 13 and 6 MYA appears not to have significantly increased the risk for schizophrenia (except possibly for some markers in the MHC region),

Our results are also in line with the findings of Srinivasan et al. (2017), who failed to find evidence of enrichment of schizophrenia using genomic markers of evolution dating back to 200 MYA. The same authors also reported enrichment of association for regions of more recent evolution in modern humans (Srinivasan et al., 2015). Interestingly, one of the evolutionary proxies used by Srinivasan and colleagues (2017), namely HARs, also showed enrichment for height, similar to our recent study (Banerjee et al., 2017). This suggests that regions controlling both genomic and epigenomic variation in height may also be driven by recent evolution. Finally, our results agree well with the observation by Srinivasan et al. (2017) of some involvement of the MHC in an early evolutionary context.

Although our results are in line with several findings in the field, the current methods have some limitations. Highly polygenic traits such as schizophrenia have a large number of genetic loci contributing to the aetiology of a disease (Bulik-Sullivan et al., 2015; Schork et al., 2016). The ability to detect these large numbers of genetic loci is dependent on the sample size and adequate statistical power (Schork et al., 2016). Consequently, the polygenic enrichment methods may be limited by the statistical power of the respective GWAS and trait polygenicity. Furthermore, in the INRICH analysis that uses LD-clumping of SNPs at *p<*10e-3 to *p<*10e-8, higher *p*-value thresholds (e.g. *p<*10e-3) still include SNPs of lower *p*-values, even though they become progressively smaller minorities. Thus, although higher *p*-values increase the number of LD-clumps tested, we do not expect this to increase the Type I error rate (Lee et al., 2012).

In conclusion, our results suggest that methylation markers tracing an evolutionary period dating back to 13 MYA (primate DMRs) are not enriched for schizophrenia markers, unlike methylation markers from a recent timeframe (non-primate DMRs) (Banerjee et al., 2017). Taken in consideration with previous studies of genomic markers of evolution dating back 200 MYA (Srinivasan et al., 2017), our results support the hypothesis that the origins of schizophrenia lie in more recent evolutionary events, possibly after the divergence of modern-day humans from Neanderthals and Denisovans.

## Appendix A. Supplementary data

### Supplementary Information

Additional Methods, Figures and Tables

### Supplementary Dataset 1

Annotation of human hypo-methylated regions with markers of schizophrenia

